# Age-related change in adult chimpanzee social network integration

**DOI:** 10.1101/2021.06.25.449973

**Authors:** Nicole Thompson González, Zarin Machanda, Emily Otali, Martin N. Muller, Drew K. Enigk, Richard Wrangham, Melissa Emery Thompson

## Abstract

**Background:** Social isolation is a key risk factor for the onset and progression of age-related disease and mortality in humans, yet older people commonly have narrowing social networks. Few models explain why human networks shrink with age, despite the risk that small networks and isolation pose. We evaluate models grounded in a life history perspective by studying social aging in wild chimpanzees, which are long-lived and show physical decline with age.

**Methodology:** We applied social network analysis to examine age-related changes in social integration in a 7+ year mixed-longitudinal dataset comprised of 38 wild adult chimpanzees (22 F, 16 M) in the Kanyawara community in the Kibale National Park, Uganda. Metrics of social integration included social attractivity and overt effort (directed degree and strength), gregariousness (undirected strength), social roles (betweenness and local transitivity), and embeddedness (eigenvector centrality) in grooming and spatial association networks.

**Results:** Males reduced overt social effort yet increased in attractivity, roles in cliques, and embeddedness. Females were overall less integrated than males, and their decreased integration with age suggested social avoidance. Effects of age were largely independent of rank. Both sexes maintained highly repeatable inter-individual differences in several aspects of integration, particularly among mixed-sex partners.

**Conclusions and implications:** As in humans, chimpanzees experience age-related declines in social effort. However, important facets of integration aged more similarly to humans in non-industrialized vs. industrialized societies, suggesting an evolutionary social mismatch between conserved declines in effort and dynamics of industrialized society. Lastly, individual and sex differences have the potential to be important mediators of successful social aging in chimpanzees, as in humans.

## Introduction

Social isolation leads to an increased risk of age-related morbidity, mortality, and cognitive decline across a number of industrialized human populations (Cohen, 2004; Holt-Lunstad et al., 2010; Umberson & Karas Montez, 2010). Equally, social ties curb the risk of mortality in a broad range of social animals (Snyder-Mackler et al., 2020; Thompson, 2019). The social ties that individuals form with partners over time and the networks in which they are integrated are important sources of support, i.e. social capital, including access to tangible help, information, and secure and stable environments (Cohen, 2004; Thompson, 2019). Despite the advantages of social integration, humans commonly shrink their network of social partners with age and reallocate social effort towards a small subset of partners (Cornwell et al., 2008; David-Barrett et al., 2016; Wrzus et al., 2013). A major goal in social gerontology has therefore been to understand the patterns that distinguish “successful” social aging from pathological aging (Cornwell et al., 2008; Rowe & Kahn, 2015). To contribute to this goal, our present study examines patterns of social aging using a mixed-longitudinal behavioral dataset from one of our closest evolutionary relatives, wild chimpanzees. Although human and chimpanzee social worlds differ, recent evidence shows that male chimpanzees exhibit striking similarities to humans in how their dyadic friendships change with age (Rosati et al., 2020). We expand on work from Rosati et al. (2020), by evaluating several life-history based drivers of social aging, and characterize multiple dimensions of sociality using a suite of social network integration measures in both males and females (Table 1 & Supplement).

**Table 1.** Guide to a) Individual network measures, where individual of interest is “ego” & b) Explanatory models of social aging tested in this study and their predicted changes in social integration.

| a) Network measure | Functional Term | Technical description |
| --- | --- | --- |
| *In - Degree Strength* | Social Attractivity | Attention received:<br>Number of partners that groom ego<br>Summed frequency of ego’s grooming received |
| *Out - Degree Strength* | Overt social effort | Attention given:<br>Number of partners that ego grooms<br>Summed duration of ego’s grooming given |
| *Strength (undirected)* | Gregariousness | Ego’s time spent in proximity ($\leq 5$ m) to a partner. |
| *Betweenness\** | Social role - Bridging | Number of shortest paths between any two network members that pass through ego |
| *Local Transitivity* | Social role – Clique member | Proportion of ego’s partner that are also partners with each other |
| *Eigenvector Centrality* | Embeddedness – influence & access to information | Individuals with high eigenvector centrality have many partners who themselves also have many partners. |
| b) Model of social aging | Predictions |  |
| *Physiological constraints* | All network measures of integration ↓ with age. |  |
| *Social selectivity* | ↓ Out-degree, same Out-strength,<br>↑ Transitivity with age. |  |
| *Social attractivity* | ↑ In-degree, In-strength, Betweenness and/or Centrality with age. |  |
| *Social status* | Dominance rank drives variation in integration with no independent effect of age.<br>Sexual status moderates any age-effect on female integration with no main effect of age. |  |
| *Individual differences* | Repeatable inter-individual differences explain significant amount of variation in integration, with or without age-effects. |  |

\*All SNA measures from Betweenness down are calculated with weighted and undirected edges.

Hypotheses for age-related declines in sociality in humans have focused on human-specific causes, such as shifts in cognitive-affective priorities with age that are driven by a perception of remaining lifetime (Carstensen et al., 1999), broken-down systems of extended family support in industrialized society (Cornwell et al., 2008), and/or significant life events that change social circles (e.g., retirement, Wrzus et al., 2013). Humans, however, are not the only animals that exhibit decreased social integration with age (e.g. red deer, Albery et al., 2021; macaques, capuchins, lemurs, reviewed in Machanda & Rosati, 2020; yellow-bellied marmots, Wey & Blumstein, 2010), and chimpanzees exhibit a suite of features associated with human social aging, including a positivity bias and strengthening of close friendships (Machanda & Rosati, 2020; Rosati et al., 2020). Thus, valid interpretations of social aging require a more generalizable framework, such as that offered by life history theory. Under such theory, individuals are predicted to use social behavior to adjust to physiological priorities and environmental challenges that vary by life stage and individual history. Key to this perspective, is that social partners are a potential source of both stress and support (Cohen, 2004; Thompson, 2019). Because of tradeoffs in the costs and benefits of sociality, older individuals’ sociality may be energetically constrained by physiological senescence and shifting reproductive priorities. Comparative studies are essential for this perspective to spread in social gerontology because they help situate human behavior and biology in its evolutionary context. Chimpanzees are a useful model of such tradeoffs in human social aging as they provide a social and physiological system that is similar to humans yet independent of advanced future-oriented cognition and contemporary human societal structures.

### Chimpanzee social network data

Chimpanzees are a tractable comparative model for human social aging, in part, because they overcome common biases in human behavioral data (Althubaiti, 2016). Holt-Lunstad et al’s (2010) important meta-analysis emphasizes the importance of structural measures of social integration (e.g. objective quantification) in predicting human morbidity and mortality, relative to functional measures (i.e. perceived experience). Data from habituated non-human primates consist of direct observations of social behavior that are suitable for constructing structural measures of social integration, including number of social ties, frequency of social contact, social roles, and overall embeddedness within networks, where each improves health outcomes and lower mortality risk in humans (Cohen, 2004; Holt-Lunstad et al., 2010). In this study, we employ social network analysis (SNA) as a powerful and standardized tool to quantify each of these structural features of individual social integration, with the advantage of incorporating direct and indirect ties that situate individuals within groups as a whole (Table 1 & Supplement).

### Study system

We used social network analysis to measure age-related changes in social integration in wild, adult chimpanzees (*Pan troglodytes*) in the Kanyawara community in the Kibale National Park, Uganda. Chimpanzees live in large communities that are closed, facilitating characterization of true global networks, and they associate in a fission-fusion pattern which allows for inter-individual variation in social integration. Although chimpanzee social life lacks important components of human social networks such as marriage, nuclear families, and a grandmothering stage of life for females (Emery Thompson, Jones, et al., 2007), chimpanzees do maintain strong ties with kin (Foerster et al., 2015; Mitani, 2009). They also have long lifespans (maximum in the wild ca. 65 years, Wood et al., 2017) and experience age-related declines in physical condition (Emery Thompson et al., 2020). Chimpanzees demonstrate stark differences in social tendencies between sexes. Males interact more frequently than females and remain in their natal communities for life, where they benefit from cooperative coalitions with other males to rise in dominance rank and access mates (Gilby et al., 2013). Females, in contrast, are less gregarious and less socially interactive than males (Wrangham, 2000), although this can vary somewhat with local ecology and community demographics (Wittiger & Boesch, 2013). Although female chimpanzees are less likely to form strong ties with one another than are males, strong female-female ties do occur (Foerster et al., 2015). Both males and females form linear dominance hierarchies that are associated with priority of access to fertile females for males (Muller et al., 2020), high quality feeding areas for females (Emery Thompson, Kahlenberg, et al., 2007), and higher reproductive success in both sexes (Emery Thompson, Kahlenberg, et al., 2007; Pusey et al., 1997; Wroblewski et al., 2009).

We evaluated male and female age-related change in social dimensions quantified by 8 social network measures (Table 1 & Supplement): social attractivity or attention received (in-degree, in-strength), overt social effort (out-degree, out-strength), gregariousness (i.e., overall time in spatial association, or proximity strength), social roles (local transitivity and betweenness), and overall embeddedness within the community (eigenvector centrality). For a full explanation of the choice of network measures, including their functions and known changes with age, see Supplement. We evaluated rates of grooming and spatial association as the currencies of the network. Because inter- and intrasexual selective pressures have differentially shaped the form and function of male-male, female-female, and male-female social relationships in chimpanzees (e.g. Gilby & Wrangham, 2008; Machanda et al., 2013), we evaluated integration within both mixed and same-sex adult networks to capture age-related changes in these functionally distinct social realms. Because social status influences both sociality and fitness, and varies with age (Braveman et al., 2011; Clutton-Brock & Huchard, 2013; Emery Thompson, Jones, et al., 2007; Muller et al., 2006), we tested and controlled for the effects of dominance rank and sexual receptivity on sociality. Lastly, we evaluated the consistency of individual differences in social traits, because personality can influence morbidity and mortality in humans and animals (Altschul et al., 2018; Cohen, 2004) and the efficacy of human social interventions (Chapman et al., 2014).

We tested changes in social network integration for consistency with 5 explanatory models (Table 1). First, under the <u>physiological constraints</u> model, the physical limitations of aging are predicted to lead to progressive social isolation, associated with decreases in all integration measures. Second, the <u>social selectivity</u> model posits that the benefits of particular ties are balanced against age-related constraints, such that social interaction is prioritized towards fewer, more valuable relationships. Under this model, we predict that individuals decrease the number of social partners they direct effort toward (lower out-degree), but that the total effort does not change (maintained out-strength). Further, under this model, partners become collectively more familiar or more cliquish with age (higher transitivity), as observed in human age-related selectivity. Third, under the <u>social attractivity</u> model, older animals attract more social partners (regardless of their dominance status), resulting in greater attention received via either more partners or increased duration of attention (higher in-degree or in-strength), and a greater likelihood of bridging and/or being embedded among network members with age (higher betweenness and/or centrality). Fourth, the <u>social status</u> model predicts that changes in sociality over the life course are specifically linked to age-associated changes in dominance rank and/or sexual status. This model predicts that aging *indirectly* influences sociality via changes in status but does not have an independent effect. Finally, we examined the potential for <u>individual</u> <u>differences</u> to shape levels of integration, alone or in combination with age effects.

## Methods

### Data Collection

Data were collected on 38 permanent residents (22 F, 16 M) of the Kanyawara Community in the Kibale National Forest, Uganda from Aug 2009 to Dec 2017 (full Data collection methods and Ethical statement in Supplement). Subjects ranged from 12 – 57 years old (Figure 1). In total, data consisted of 3371 focal follows, with subjects observed as focals for 133 ± 73 hours per year (mean ± sd) and as party members during focals for 1033 ± 588 hours per year.

### Analysis

We used the R package igraph v. 1.2.6 to create network graphs and measure individual-level network integration in 4 types of annual networks: networks based on grooming or spatial association within 5 m (proximity) and among members of both sexes (mixed-sex) or of the same sex (i.e. all male, all female; Supplement). We calculated **in-degree, in-strength, out-degree,** and **out-strength** for directed grooming networks; undirected **strength** in proximity networks; and **local transitivity**, **betweenness**, and **eigenvector centrality** in both total undirected grooming and proximity networks. Although grooming and spatial association behavior are similar in their affiliative and tolerant tone, each integration measure from one network behavior type was not on average correlated with the same measure from the other, within indviduals observed ≥ 3 years (N = 30, range average Spearman’s rhos −0.10 – 0.51, all p ≥ 0.39). All measures apart from in-degree and out-degree were weighted in an effort to capture variation in both number of social partners and frequency of social interaction. We did not calculate individual degree in proximity networks (i.e. an individual’s unweighted number of annual spatial associates) as such networks were often fully connected on an annual basis.

To evaluate changes in network integration with age, we constructed general additive mixed models (GAMMs) in the R package mgcv v. 1.8-31 (S. N. Wood, 2017). General additive models were useful for our age analysis because we expected social integration to vary over the life course in a non-linear fashion, as reproductive priorities and physiological constraints demonstrate non-monotonic changes with age. The curviness of non-linear relationships in GAMMs (smooths) are determined by the number of basis functions for each fixed effect, optimized for each model and effect (with mgcv::gam.check), All smooth parameters were estimated with restricted maximum likelihood. Each network integration measure was modeled as a response with either a Gaussian or Gamma error distribution and a log-link function, based on model diagnostics with the mgcv::gam.check function. We ran our models in two sets to evaluate age effects independent of social and reproductive status (Table 2). In both sets, we included age as a smooth term (age calculation in Supplement), estimated by thin plate splines with a k of 5 optimized by the mgcv::gam.check function, and individual ID as a smoothed random intercept. In set 1, we included annual dominance ranks based on aggressive interactions (calculation in Supplement) for both males and females in mixed and same sex networks. In set 2, we included annual time swollen (calculation in Supplement) for females’ alone in mixed sex networks. In time swollen models, we included an interaction between female age and time swollen, as we expected females in estrus to be more attractive to males when they were older (Muller et al. 2006). We lastly included an analysis of models with age alone as a predictor (results in Tables S9-13) for readers interested in the unconditional effect of age on integration measures.

**Table 2.** GAMM compositions: testing effects of age on social integration independent of annual dominance rank and time swollen.^†^

| Approach | Network composition | Network behavior | Responses | Linear Predictors and Smooth Terms |
| --- | --- | --- | --- | --- |
| Rank-independent age effects | Mixed-sex | Grooming & ≤ 5 m Proximity | In-Degree, Out-degree*, In-Strength, Out-Strength, Strength, Local Transitivity, Betweenness, Eigenvector centrality | Sex + s(Age, by = Sex, k = 5) + s(Rank, by = Sex, k = 5) |
|  | Same-sex | Grooming & ≤ 5 m Proximity | “ ” | s(Age, k = 5) + s(Rank, k = 5) |
| Time swollen-independent age effects (females only) | Mixed-sex | Grooming & ≤ 5 m Proximity | “ ” | s(Age, k = 5) + s(Rank, k = 5) + s(Time swollen, k = 5) + ti(Age, Time swollen, k = 5) |

<sup>†</sup> All models included individual ID as a random effect: s(ID, bs = “re”)
\*In-Degree and Out-Degree calculated based on directed grooming networks, other measures on undirected networks.

Generalized additive models as implemented by the mgcv package are robust to concurvity (Wood, 2017), an issue similar to collinearity but for non-linear models. Thus, although male and female dominance rank, and female annual time swollen, were strongly related to age (Table S1), estimates of their independent effects on integration were stable. Permutation methods were used for significance testing of the influence of predictors on integration measures (Supplement). This method, where effect sizes are compared to those from models run on node-randomized permutations of observed data, reduces the risk of type I error that typically grows with multiple testing, and so avoids the need for correction of multiple comparisons (Farine & Whitehead, 2015). Consistent inter-individual differences in social integration (repeatability) were evaluated by variance decomposition of each GAMM’s random effect of individual ID, identical to methods employed in linear models (Nakagawa et al., 2017) and their significance calculated via permutation methods used in models of social aging. (Supplement).

## Results

Age-related changes in social integration measures for both males and females overwhelmingly occurred in grooming rather than proximity networks (Table 3). We therefore focus on age-related changes in grooming networks in our presentation of results and their discussion.

**Table 3.**
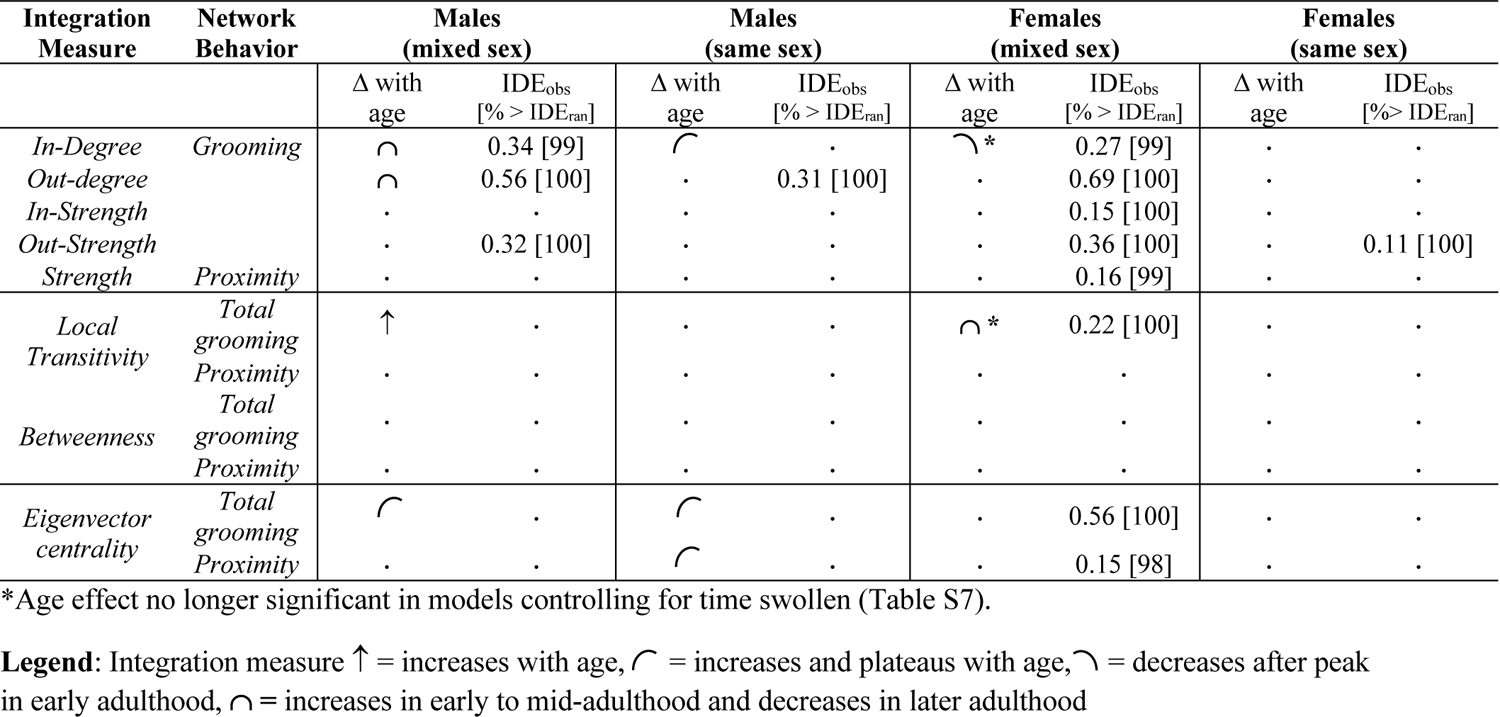
Summary of results: Age-related changes in social network integration independent of dominance rank. Shape and arrows describe significant relationships between age and a given network measure (see Legend; full model results in Tables S3-8). Dots indicate a non-significant relationship with age. Significant repeatability of integration measures given as IDE_obs_ (observed deviance explained by individual ID in GAMM). Significance of IDE_obs_ was evaluated by the proportion of 1000 deviances explained by ID in GAMMs on node-randomized data (IDE_ran_) that IDE_obs_ was less than (full Table S8).

### Males

Across analyses, male chimpanzees exhibited three notable areas of changes in integration with age (Table 3 & S3-6, Fig. 2 & S1). First, age significantly affected the number of partners males groomed with (in/out-degree), but not their time spent grooming (in/out-strength, Table 3 & S3). Older males declined in the number of mixed-sex partners that they gave and received grooming from (out & in-degree), with males grooming with the most partners of either sex in their late 20s (Fig. 2). Although this might suggest an influence of dominance rank on male sociality, which also shows a concave relationship with age, these effects were independent of rank (Table 3 & S3). In contrast, males received grooming from the most male partners in their 30s and 40s (in-degree, Fig. 2), and while this declined somewhat amongst the oldest males, they still received grooming from more partners than did the youngest adults. Age only predicted a decrease in the number of partners males groomed with (out & in-degree) in mixed-sex networks (Table 3 & S3-4), indicating that aging led males to groom with fewer females, rather than males. Second, males’ grooming partners in mixed-sex networks were more likely to groom one another as males aged (linear increase in local transitivity, Fig. 2), indicating that their reduction in grooming partners (out & in-degree) was accompanied by an increased ‘cliquishness’ with age (Table 3 & S3). Third, males’ embeddedness among partners (eigenvector centrality) changed with age in all networks examined, apart from mixed-sex proximity. In each network, older males were more central than younger males, usually after declining somewhat from their peak centrality in mid-adulthood (Table 3 & S3,4, & 6, Fig. 2 & S1). The only instance in which male dominance rank had an effect on integration in the absence of age was males’ linear increase in centrality with rank in mixed-sex proximity networks (Table S5, Fig. S3). Males also maintained highly repeatable inter-individual differences in overt social effort (out-degree and out-strength) and their attractivity (in-degree and in-strength), particularly among mixed sex partners (Table 3 & S8).

### Females

Relative to males, females displayed low levels of integration and few age-related changes in network measures (Table 3, Fig.2, direct sex comparisons in Supplemental Results & Tables S3 & S5). Those rare instances of age-related change were typically declines. Females received grooming from fewer partners with age (in-degree, Table 3 & S3, Fig. 2) and, in contrast to males, females’ grooming partners were less likely to groom one another with age in mixed-sex but not same-sex networks (reduced grooming transitivity, Table 3 & S3-4, Fig. 2). These declines with age signaled that females were grooming with fewer males, mirroring the same pattern in male transitivity.

After controlling for female’s annual time swollen, age no longer had any independent effect on female social integration in mixed-sex networks (grooming in-degree, local transitivity) although time swollen was not significantly related to either measure (Table 3 and S6). Annual time swollen did, however, independently decrease grooming out-strength (Table S6A) and interacted with age such that older females received more grooming (in-strength) and were more central in proximity networks with more annual time fully swollen (Table S6B, Fig. S4a & b).

The single instance in which female dominance rank influenced integration, without an independent effect of age, was a linear increase in time spent grooming fellow females with increases in rank (out-strength, Table S4, Fig. S4). Females showed repeatable inter-individual differences in all measures among mixed-sex partners except betweenness and local transitivity in proximity networks (Table 3 & S8). Among all-female partners, females were repeatable only in the time they spent grooming other females (out-strength).

**Table 4.** Summary of evidence consistent and inconsistent with 5 models of social aging.

| Model of social aging | Evidence consistent with model in bold, inconsistent unbolded |  |
| --- | --- | --- |
|  | Male | Female |
| *Physiological constraints* | **↓ In-Degree<sup>MS</sup>** **↓ Out-Degree<sup>MS</sup>** | **↓ In-Degree<sup>MS</sup>** , **↓ Transitivity<sup>MS</sup>** |
| *Social selectivity* | **↓ Out-Degree<sup>MS</sup>**, same Out-Strength,<br>**↑ grooming Transitivity<sup>MS</sup>** |  |
| *Social attractivity* | **↑ In-Degree<sup>SS</sup>** and **↑ grooming and proximity Centrality** | **↓ In-Degree<sup>MS</sup>** |
| *Social status* | Multitude of age-related changes in integration are independent of rank.<br>**↑ proximity Centrality with rank & no age effect.** | **↑ Out-Strength<sup>SS</sup> with rank & no age effect.**<br>**↑ proximity Centrality<sup>MS</sup> and grooming In-Strength<sup>MS</sup> with time swollen when older & no main effect of age.** |
| *Individual differences* | **Measures of social attractivity<sup>MS</sup> and overt social effort repeatable** | **Majority of network measures<sup>MS</sup> are highly repeatable.** |

<sup>MS</sup> change occurs in mixed-sex networks only
<sup>SS</sup> change occurs in same-sex networks only

## Discussion

In this study, we analyzed age-related changes in key dimensions of social integration (social attractivity, overt effort, gregariousness, social roles, and embeddedness) in wild chimpanzees, to evaluate 5 explanatory models of social aging: physiological constraints, social selectivity, social attractivity, changing social status, and individual effects. Our results indicate that aging influences sociality in both direct and indirect ways, but that these influences differ between the sexes. We further find that overt social behavior, such as grooming, is a primary way that chimpanzee social integration varies with age, whereas spatial association in close proximity is less informative. Overall, our results argue against a simple physiological constraints or social status-dependent model for social aging in chimpanzees and suggest that male social integration, in particular, is more dependent on age than rank. Additionally, our data provided evidence of individually-stable social phenotypes in both males and females, suggesting that like humans, individual chimpanzees may be predisposed to more or less successful aging trajectories (Rowe & Kahn, 2015). Here, we discuss patterns of male and female social aging separately in light of our 5 explanatory models and consider the implications of these patterns for human social aging and age-related disease.

### Males’ age-related changes in integration

Male patterns of social integration were broadly consistent with both social selectivity and attractivity models of social aging, which posited an age-related focus on valuable social ties and increases in attention received and embeddedness, respectively. Older males focused grooming on a small set of partners that were increasingly connected with one another (lower in & out-degree, maintained strength, higher transitivity, Fig. 2). Their selective focus parallels other findings from this field site using different measures of sociality, where males formed more equitable relationships with one another as they aged (Rosati et al., 2020). However, in this analysis, the effects of aging on cliquishness (grooming transitivity) and overt social effort (out-degree) were most affected by decreased interactions with females, as these two dimensions changed in mixed-sex but not in all-male networks. Kanyawara males’ selectivity does not result from a narrow focus on kin, as few close kin pairs exist in our dataset. Though it is likely that chimpanzees do not have knowledge of their impending mortality (a central feature of one major theory of human social aging, Carstensen et al., 1999), aging male chimpanzees may nevertheless shift their social goals with age. For example, males’ strong increase in grooming cliquishness (transitivity) may reflect a preference for predictability and stability that increases with age. Further, young male chimpanzees cultivated a diversity of both male and female grooming partners (in & out-degree) (Fig. 2 & S1), indicating motivation to secure allies and affiliate with potential mates as young adults (Enigk et al., 2020), which is consistent with ‘information gathering’ goals (Carstensen et al., 1999).

Male social patterns also indicated that age *per se* increased male attractivity, as older males received grooming from more male partners (in-degree), were more cliquish (grooming local transitivity), and were more embedded within the community than younger males (grooming and proximity centrality) independently of dominance rank (Table 3, Fig. 2 & S1). Older male chimpanzees exhibit declining physical condition (Emery Thompson et al. 2020), which emphasizes that an older male’s value as a social partner lies in reasons other than physical ability or rank-based benefits. Studies of other non-humans suggest that older individuals are valued social partners due to their accumulated knowledge and experience (reviewed in Brent et al., 2015). For chimpanzees, while it is possible that older males have increased ecological knowledge that is of value to others, there is no direct evidence of this, and it is not clear that grooming relationships would be necessary to benefit from such knowledge. Instead, is it more plausible that older males’ have social and political experience that can assist younger, less experienced partners to navigate competitive environments. Further, older males exhibit less aggression (Muller et al., 2020), and tolerance is a potentially important factor in their attractivity and the transmission of knowledge (Thornton & Clutton-Brock, 2011). Indeed, older male chimpanzees have higher siring success than would be predicted by their ranks and aggressive tendencies (Muller et al., 2020), one likely pay-off of knowledge and cooperative ties (Gilby et al., 2013). Male chimpanzees’ maintenance of high embeddedness in old age was similar to social patterns in the socially dominant sex in other primates (Machanda & Rosati, 2020).

### Females’ age-related changes in integration

Female social integration was consistently low relative to males’ and, in their old age, females appeared to neither groom nor maintain proximity with any adult partners (Fig. 2 & S1). They were highest in the number of partners that groomed them (in-degree) and the cliquishness of their grooming partners (local transitivity) in their late teens and 20’s but declined thereafter (Fig. 2). That age-related changes were exclusively declines paints a picture of older female chimpanzees’ withdrawal from adult social partners. It is unlikely that females, but not males, were constrained purely by physical senescence, given that older males show more pronounced effects of declining physical condition than do older females (Emery Thompson et al., 2020). Instead, age-related aspects of their reproductive and social status appeared to shape female social integration.

One source of females’ declining integration was decreased interactions with males. Although our analyses attempted to control for mating interactions as a driving social force, we found that annual time swollen did influence certain relationships between female integration and age, suggesting that changes in other reproductive factors, such as sexual attractiveness or the presence of dependent offspring (Otali & Gilchrist, 2006), could alter affiliative relationships with males. Older females are more desirable mating partners for males (Muller et al., 2006), as evidenced in this study by their increased grooming received and proximity centrality when sexually swollen (Fig. S4a & b), and this puts them at increased risk of sexual coercion (Muller et al., 2007). Reducing interactions with males overall may thus be a strategy to reduce coercion (Wrangham, 2002). Alternatively, avoiding males could circumvent the particularly high feeding competition that associating with males imposes (Emery Thompson et al., 2014). Indeed, although socializing offspring can bring females into association (Lehmann & Boesch, 2009; Murray et al., 2014), energetically demanding states such as lactation lead females to avoid social foraging (Otali & Gilchrist, 2006) and to spend considerable amounts of time alone (Lee et al., 2021). Additionally, younger females who have newly immigrated to a community use affiliation with males to protect them from other females, but they reduce these affiliations once they are established in the community and begin to rise in status (Kahlenberg, Emery Thompson, et al., 2008; Kahlenberg, Thompson, et al., 2008). Such underlying drivers of fewer interactions with males suggest that females’ declines in integration with age stem from social avoidance, a form of reduced effort.

Females’ social status was a lone predictor explaining their social effort towards fellow females (out-strength). Female dominance rank at Kanyawara increases with age (Kahlenberg, Emery Thompson, et al., 2008), as at other sites (Foerster et al., 2016). Although females appeared to decrease overt social effort towards fellow females with age (Fig. 2) they in fact invested more time in female partners as they became higher-ranking. This effect of rank contrasts somewhat with that expected in female-philopatric species, where high-ranking females often maintain more geographically central positions among group members (Kalbitzer et al., 2017) and receive more grooming than low-ranking females (Schino, 2001). In this study, high-ranking female chimpanzees groomed other females more but were no more socially central and did not receive more grooming than low-ranking females. High-ranking females tend to inhabit higher quality core areas in Kanyawara (Kahlenberg, Emery Thompson, et al., 2008), and such access to resources may free females from either energetic or foraging-related time constraints on social interaction. Additionally, young females are subject to harassment from older females (Emery Thompson et al., 2010; Kahlenberg, Thompson, et al., 2008), thus higher rank conferred by age may simply allow females the power and confidence to associate more freely, with fewer concerns of aggressive competition. In either case, the result highlights a peculiar feature of female chimpanzee social life, in which same-sex sociality is constrained by competition. Although the effects of social status on female integration covaries, on average, with female age, they are not explained by aging, *per se*.

### Significance of individual effects on integration

Kanyawara chimpanzees maintained stable between-individual differences in several dimensions of social integration (Table 3), i.e. certain chimpanzees were, for example, consistently more gregarious or embedded than others, similar to chimpanzees in the Taï Forest, Côte d’Ivoire (Tkaczynski et al., 2020). Thus, if social integration is important to health in chimpanzees, as it is in humans and many other species, individuals’ social phenotypes could be more or less conducive to successful aging (Rowe & Kahn, 2015). In other species, such individual variation facilitates roles in cooperation (Bergmüller & Taborsky, 2010) and, in male chimpanzees, may be involved in alternative strategies to achieve dominance (Foster et al., 2009). As individual differences explained more variation in female social integration than did rank or age, further examination of the attributes driving female chimpanzees’ differences in social integration is well warranted.

### Comparisons to and implications for human social aging

Several patterns of social aging in chimpanzees were consistent with those in industrialized human populations, but others diverged in important ways. Like industrialized humans, both male and female chimpanzees at Kanyawara increased their number of social partners in early and mid-adulthood and declined thereafter (David-Barrett et al., 2016; Fung et al., 2001; Wrzus et al., 2013). Further, male chimpanzees participated in tighter social cliques with age, rather than bridging otherwise unconnected partners, like many men (Cornwell et al., 2009). However, unlike most men in industrialized societies, chimpanzee males sustained high overall levels of integration into old age, with high attention received (in-degree) and embeddedness (centrality). Relatedly, chimpanzees’ sex differences in social aging were largely opposite to that observed in industrialized populations, where women consistently have larger networks than men after early adulthood (Bhattacharya et al., 2016; Cornwell et al., 2008). Further, there are no obvious sex differences in social selectivity with age in studied humans (Carstensen et al., 1999), but chimpanzee males appeared to be more socially selective with age than females, given males’ overall higher rates of integration and increased cliquishness with age (Fig. 2).

Where Kanyawara chimpanzees contrasted with industrialized humans, their sociality appeared to age more similarly to humans in non-industrialized settings, where social networks are primarily based within small communities. Although data on social aging from non-industrialized societies are admittedly sparse and preclude indisputable comparisons, several similarities are apparent. Men in non-industrialized societies, such as in Tsimane forager-horticulturalists and Nyangatom agro-pastoralists, often retain significant prestige even in old age, similar to male chimpanzees (Glowacki & von Rueden, 2015). Further, female chimpanzees’ low social integration relative to males resembles the situation of women in some patrilocal and non-industrialized societies that disperse at marriage and are limited in replacing kin relationships with new non-kin partners (Scelza, 2011; Wood & Eagly, 2002). For example, in Himba semi-nomadic pastoralists, women are often hindered in their travel to visit kin for social support because of mate-guarding within their marriage (Scelza, 2011). Among the Tsimane and nomadic Saami, women also face trade-offs between having large, cooperative social networks and attending to duties of intra-household labor and childcare (Anderson, 1983; von Rueden et al., 2018). In each case, women are socially limited by male reproductive tactics and their reproductive priorities, similar to female chimpanzees. Comparing social aging in this community of chimpanzees with future studies on age-related changes in sociality in diverse human cultures, other chimpanzee communities, and other closely-related apes, would allow even greater inferences into how ecological variability in gender roles shapes social aging, and into the nature of humans’ ancestral social environments.

Similarities in non-human primate and human social aging suggest their similar and potentially evolutionarily conserved drivers. Given that chimpanzees’ and other primates’ likely lack abstract knowledge of their impending mortality, their decreasing sociality likely results from the constraints of variable costs of social interaction, and their selectivity likely functions to maintain the most beneficial of social ties. Sex-specific patterns of social aging in this study emphasize that physiological priorities drive social decision-making.

### Implications for human age-related disease

Although social integration is well-linked to fitness in non-human primates (Snyder-Mackler et al., 2020; Thompson, 2019), whether social integration moderates age-related declines in physical health in non-human primates is currently an open question. Although we did not yet test these effects here, we hypothesize that chimpanzees’ and humans’ shared tendencies to decrease social effort and become more socially selective with age are not in themselves evidence of pathology. Instead, they may have been adaptive strategies for coping with the constraints of aging in past social environments that are now disadvantageous in industrialized society (Gurven & Lieberman, 2020).

In the evolutionarily novel environment of industrialized nations, humans’ conserved tendencies to decrease social effort and increase selectivity may be at particular risk of developing into isolation, with strong physical consequences. In terms of physiology, advanced physical and mental deterioration during humans’ extended lifespans could make the effects of decreased integration on physiological function particularly dramatic. In terms of culture, many industrialized societies lack deference to older people (North & Fiske, 2015) and cohesive communities that endure for a lifetime (Höllinger & Haller, 1990). In contrast, chimpanzees experience a relatively permanent social community, and this alone could preserve older male chimpanzees’ network size and attention received, and older females’ social status. Similarities in social aging between chimpanzees and people in non-industrialized societies reinforces the likelihood that industrialized humans have recently departed from social settings in which community stability is a norm and social isolation unlikely. Again, greater research on social aging in a diversity of non-industrialized societies can further elucidate and reinforce reference points of successful social aging and vulnerabilities to related diseases. Such insights can inspire and support the rationales of certain social interventions for older people, such as prioritizing stability and control in older adults’ social environments over a manufactured sense of belonging or introduction of new social ties (Cohen, 2004; Fung et al., 2001; Umberson & Karas Montez, 2010).

## Acknowledgements

We are grateful to the staff and field assistants of the Kibale Chimpanzee Project for their efforts in collecting all behavioral data. We acknowledge and give thanks to Stephanie Fox, Kathrine Starkweather, Kris Sabbi, and Shauhin Alavi for conversations that improved the interpretation of our analyses. We also thank the Uganda Wildlife Association, Makerere University Biological Field Station, and the Uganda National Council for Science and Technology for their support and permission to conduct research in Kibale National Park.

## Data Availability Statement

Relevant data and scripts for analysis are publicly available in author NTG’s GitHub page at https://github.com/Gavago/Social-aging-in-wild-adult-chimpanzees.

## Supplemental Background

### Justification of Social Network Measures: functions and changes with age

Social network analysis has the distinct advantage of providing individual measures of integration based on either direct or indirect ties, with the latter situating individuals within groups as a whole (Table 1). The overall number of direct social partners an individual has (i.e., **degree** centrality) represents its range or flexibility in possible sources of social support and resources (Donald & Ware, 1984; Thompson, 2019). Greater frequency of contact or association with partners (i.e., **strength** or intensity of social ties), indicates individual gregariousness and the presence of preferential relationships that can predict reliable support (Bray & Gilby, 2020; Granovetter, 1983; James, 2000; Mitani, 2009; Young et al., 2014). In humans, although degree generally decreases with age (Cornwell et al., 2008; David-Barrett et al., 2016; English & Carstensen, 2014; Fung et al., 2001; Wrzus et al., 2013), strength does not always follow the same pattern, sometimes decreasing and sometimes remaining the same, indicating a relative increase among a smaller set of social partners (Carstensen, 1992; Cornwell et al., 2008).

Directional measures of degree/strength further tease apart overt forms of individual social attractivity vs. social effort, or attention received vs. given. In Barbary macaques, for example, adult females maintain the same number of groomers and amount of grooming received as they age (**in-degree** and **in-strength**), but reduce their overt social effort by grooming fewer individuals less often (out-degree and out-strength, Almeling et al., 2016). Across animals, both social attractivity and effort change with age. For example, older individuals sometimes attract more attention because of their experience, including greater political knowledge (men, Glowacki & von Rueden, 2015; von Rueden et al., 2008), ecological knowledge (female orcas, elephants, and bonobos Brent et al., 2015; McComb et al., 2001, 2011; Tokuyama & Furuichi, 2017), or reproductive parity (female chimpanzees, Anderson, 1986; Muller et al., 2006). Social effort, on the other hand, often decreases with age in many primates (reviewed in Machanda & Rosati, 2020), possibly because older and senescing individuals are simply less able to physically compete, a direct cost of sociality (Emery Thompson et al., 2020; Silk, 2007).

In humans, social roles are positions held within a group that involve both direct and indirect group ties. Roles in humans are thought to promote health by increasing one’s sense of identity and purpose (Cornwell et al., 2008; Holt-Lunstad et al., 2010) and potentially mirror several aspects of animal social behavior that similarly promote homeostasis and environmental stability (Matthews & Tye, 2019). In SNA, one measure of social role is participation in cliques, i.e. when one’s contacts interact with one another (**local transitivity**, Table 1). When social contacts form cliques it increases the likelihood that cooperation and reciprocity will ensue (Sosa et al., 2020), creating secure environments where information can be triangulated and where resources such as food and vigilance can be pooled (Cornwell et al., 2008; Hanneman & Riddle, 2005). A second measure of social role, and one often inversely related to transitivity, is an individual’s ability to bridge disparate cliques or otherwise unconnected individuals (**betweenness centrality**; Cornwell et al., 2009; Hanneman & Riddle, 2005). The benefit of bridging otherwise unconnected individuals is to uniquely access and broker information and/or to have access to distinct pools of resources (Brent, 2015; Keating et al., 2005). In dolphins (*Tursiops spp.*), for example, highly ‘between’ individuals possess greater ecological knowledge and are key in facilitating cohesion (Lusseau & Newman, 2004), and decision-making in communities (Lusseau, 2007). No human or non-human animal studies have yet examined age-related variation in social roles measured as local transitivity or betweenness *per se*. However, people’s increased participation in religious and volunteer organizations and focus on few, close social contacts in late adulthood suggests that humans do increase in local transitivity with age (Bhattacharya et al., 2016; Carstensen et al., 1999; Wrzus et al., 2013). Limited research indicates that humans have little to no tendencies to bridge different partners in old age (Cornwell et al., 2009; Wen Yuan et al., 2017).

Lastly, social “embeddedness” is a fundamental concept in the social determinants of health literature, highlighting that individuals derive social capital from their position within a global network of indirect ties, or “friends of friends”, including access to information and social norms (Carstensen et al., 1999; Coleman, 1988; Cornwell et al., 2008; Keating et al., 2005; Stowe & Cooney, 2015). Although widely referenced (e.g. Coleman, 1988; Granovetter, 1985), embeddedness *per se* is rarely quantified in human health studies, but can be well captured in SNA as **eigenvector centrality** (Andersen, 2013; hereafter, centrality, Table 1). High measures of centrality derive from an individual’s many and strong social ties and those of their direct contacts (Sosa et al., 2020). In non-human animals, centrality corresponds with greater food discovery (Paridae songbirds, Aplin et al., 2012), and has been shown to decrease with age in female yellow-bellied marmots (Blumstein et al., 2018), and in some primates (Barbary macaques, Rathke & Fischer, 2021) but not all those examined (rhesus macaques, Liao et al., 2018). In some species, embeddedness corresponds with decreased parasites and infection (Balasubramaniam et al., 2016; Duboscq et al., 2016), however, under some circumstances it can lead to greater pathogen exposure (Nunn, 2012; Page et al., 2017). In humans, embeddedness is thought to decline with age alongside shrinking social networks (Cornwell et al., 2008).

## Supplemental Methods

### Ethical statement

The Institutional Animal Care and Use Committees of Harvard University and the University of New Mexico approved of this study’s data collection protocol. All research was conducted in compliance with Ugandan law, with research permissions granted by the Uganda Wildlife Authority, Uganda National Council for Science and Technology, and Makerere University Biological Field Station.

### Data collection

The Kanyawara community of wild chimpanzees lives in the northern part of Kibale National Park, Uganda. From August 2009 to December 2017, pairs of field assistants of the Kibale Chimpanzee Project conducted focal follows of individual chimpanzees, wherein they attempted to follow the same chimpanzee (and that chimpanzee’s associates) through the entire active period from waking to nesting (mean ± sd = 9.8 ± 2.7 hrs per follow, N = 3371 follows). Focals were selected based on which individuals were located on a given day, prioritizing those who had been followed less recently or less frequently. If a focal was lost, another was chosen, if possible, to finish the observation day. One observer collected party composition data (all individuals within 50 m of any other) via instantaneous scan sampling every 15 minutes, while a second recorded the focal individual’s activity (e.g., resting, grooming, feeding) each minute and recorded all individuals within 5 m of the focal every 15 minutes. The average chimpanzee was a focal subject for 133 ± 73 hours per year (130 ± 78 F, 138 ± 63 M) and a party member for 1033 ± 588 hours per year (937 ± 531 F, 1184 ± 642 M; annual values Table S1). Importantly, within subjects, no annual measure of social integration in any network was, on average, correlated with annual observation time as a focal or party member (subjects observed ≥ 3 years N = 30, range of average Spearman’s rho for within-individual correlations −0.30 – 0.55, all p > 0.22).

The study examined social integration in the 22 female and 16 male adults that permanently resided in the Kanyawara community between 2009 to 2017, for a total of 200 unique chimp-years. Networks were calculated on an annual basis, but because focal data collection started late in 2009, we combined data from 2009 and 2010. Social networks included only adult individuals, including males ≥ 15 years and females ≥ 12 years. Members ranged from 12 – 57 years old, with an average age of 26.5 +/- 11.6 years (mean +/- sd), and each member contributed to 1 – 8 years of networks, with an average 5.26 +/- 2.7 years (Fig. 1). Individuals were included as annual network members if present in the community for ≥ 6 months of the year (where absence was related to their pre-immigration status or death), and if observed either > 50 hours as a focal or > 100 hours as a party member during focals. These criteria led us to omit only 15 insufficient chimp-years, resulting in full adult networks that ranged from 22 to 27 individuals, male networks from 8 to 11 individuals, and female networks from 14 to 17 individuals.

### Calculation of covariates: annual age, dominance rank, and time swollen

We calculated two dyadic indices based on grooming and proximity. Each were calculated by summing the number of focal point samples throughout the calendar year when the dyad members were observed grooming or within 5 m of one another. We note that these two measures are not mutually exclusive, as grooming partners were also recorded as within 5 m of a focal. We then controlled for the dyad members’ opportunity to associate by dividing this sum by the number of point samples in which the two were seen in the same party and one was a focal (as in Machanda et al., 2013).

We measured annual **age** at the mid-year (July 1) for all subjects. Birthdates of natal community members born after 1987 were known to within one year. Birthdates of individuals born before 1987 (most first encountered in 1983) were estimated based on body size, if immature, or by signs of relative aging, including body hair and presence of dependent offspring (see Muller & Wrangham, 2014). Immigrant, nulliparous females were assigned an age of 13, the average age when natal females are seen to disperse from the community. To calculate individual annual dominance **rank,** we averaged daily dominance ranks within sex-specific dominance hierarchies across one year. Daily dominance ranks were based on Elo ratings informed by decided agonistic interactions, as described in Emery Thompson *et al*. (2020), and standardized relative to number of individuals in the hierarchy (1 = highest rank, 0 = lowest rank). Lastly, to control for changes in reproductive activity with age, we calculated the proportion of observation days in a given year that a female was seen with a maximally tumescent swelling (**time swollen**). Mating primarily occurs when females are in this state (Muller & Wrangham, 2004), and associations with males consequently increase.

### Assessing significant changes in integration with age in GAMM models

To control for dyadic non-independence in network data, we tested the significance of patterns of social integration related to age, sex, rank, and reproductive status in GAMM models by creating 1000 randomized versions of each network, where node attributes such as sex, age, rank, time swollen (among females alone), and ID were assigned randomly within years (Farine, 2017). Node randomization preserved, and thus controlled for, annual variation in network size, sex and age composition, and potential stability in individual social tendencies. We ran our original models on these randomized data sets 1000 times each and extracted the estimated F statistics of the smooths of interest (e.g. age, rank, time swollen, age * time swollen) and linear coefficients of the categorical predictor “sex”. We then calculated the proportion of randomized F statistics and linear coefficients that fell below the observed models’ F statistic and coefficient, where proportions > 0.95 indicated a significant pattern in the smooth term and > 0.95 and < 0.05 indicated a significantly positive or negative effect of the categorical predictor.

### Calculating repeatable inter-individual differences

To evaluate the individual differences model of social aging, we measured the consistency of individual differences (i.e. **repeatability**) in each social integration measure. We calculated a repeatability statistic by partitioning the deviance explained by individual intercept (ID) in each GAMM, following methods for generalized linear models (Nakagawa et al., 2017; Schielzeth & Nakagawa, 2020). In this approach, deviance explained is used as a coefficient of variation, similar to the R^2^ in linear models, that is generalized and appropriate for GAMs (Wood, 2017). We evaluated the significance of the repeatability statistic by comparing the observed deviance explained by individual ID to 1000 deviances explained by ID in models of node-randomized data, i.e. data with randomized attributes of rank, time swollen, and ID, within years, network behavior, network type, and individual sex. An integration measure was significantly repeatable if its repeatability statistic was ≥ 95% of its random statistics. Because of large sex differences in social tendencies, we modeled male and female repeatability separately, and controlled for annual rank, and annual time swollen (for females only in mixed sex networks) as fixed effects. Significantly repeatable inter-individual differences in integration in the absence of age effects in GAMMs would indicate variation in integration resulting primarily from individual traits, whereas repeatable differences in combination with an age effect on integration would represent differences in the extent of individual integration (intercept) within an overall age-related pattern.

## Supplemental Results

### Average sex differences in integration measures

Among partners of both sexes (mixed-sex networks), males were more socially integrated than females according to all measures of grooming except for in-strength (i.e., in-degree, out-degree, out-strength, local transitivity, betweenness, and centrality; Fig. 2 & Table S3). Males also spent more time than females in association and embedded among proximity partners (higher strength and centrality, Fig. S1 & Table S5). In proximity networks with mixed-sex dyads, sexes did not differ in their tendency to form cliques or bridge otherwise unconnected partners (local transitivity and betweenness, Fig. S1, Table S5).

**Table S1.** Average annual observation times per subject as focal or party member during focal follows.

| Year | Sex | Mean ± sd focal hours per subject | Mean ± sd party hours per subject |
| --- | --- | --- | --- |
| 2010* | F | 133.9 ± 118.9 | 922.6 ± 489.9 |
| 2010* | M | 153.2 ± 50.3 | 1248.3 ± 411.7 |
| 2011 | F | 66.3 ± 40.4 | 559.5 ± 169.7 |
| 2011 | M | 70 ± 17.7 | 752 ± 170.4 |
| 2012 | F | 85.5 ± 68.6 | 481.3 ± 181.4 |
| 2012 | M | 96.5 ± 37.1 | 504.1 ± 169.7 |
| 2013 | F | 116.5 ± 48.1 | 779.4 ± 219.4 |
| 2013 | M | 116.3 ± 44.2 | 911.4 ± 196.5 |
| 2014 | F | 137.4 ± 64.6 | 699.5 ± 216.3 |
| 2014 | M | 102.3 ± 45.1 | 743.8 ± 232.3 |
| 2015 | F | 160.9 ± 59.4 | 1808.3 ± 534.3 |
| 2015 | M | 180.5 ± 54.7 | 2289.7 ± 581.3 |
| 2016 | F | 186.8 ± 76.8 | 984.9 ± 429.3 |
| 2016 | M | 219.4 ± 48.6 | 1598.7 ± 424.7 |
| 2017 | F | 152.3 ± 64.8 | 1233.1 ± 391.1 |
| 2017 | M | 175.4 ± 37.3 | 1634.1 ± 455.4 |

\*2010 = Aug-Dec 2009 & all 2010 combined

**Table S2.** Significant relationships in GAMM models between male and female age and annual dominance rank (Elo scores) and female age and annual time swollen (N females = 22 individuals, 122 female-years; N males = 16 individuals, 78 male-years). Significance evaluated with model P values. Male rank showed a concurve pattern with age. Female rank a rise and plateau with age. Female time swollen decreased linearly with age.

| Response | Predictor | F | P value |
| --- | --- | --- | --- |
| Annual dominance rank | female age | 29.2 | < 0.001 |
|  | male age | 60.2 | < 0.001 |
| Annual time swollen | female age | 19.4 | < 0.001 |

**Table S3.** GAMM models for all integration measures in <u>mixed-sex grooming</u> networks. Significant effects in bold with *. DE = total model deviance explained. Significance of the categorical variable sex evaluated with linear ß estimates, and all smooth terms (age & rank) evaluated with observed F statistics, each compared to ßs and F statistics drawn from randomized networks.

| Response | Network sex | Behavior | DE | Predictors | F<sub>obs</sub> of smooths | $\beta_{obs}$ of sex(M) | % F<sub>obs</sub> > F<sub>ran</sub> | % $\beta_{obs}$ > $\beta_{ran}$ |
| --- | --- | --- | --- | --- | --- | --- | --- | --- |
| In-Degree | mixed | Grooming | 0.81 | **sex(M)**<br>female age<br>male age<br>female rank<br>male rank | 4.4<br>6.2<br>3.59<br>5.01 | 0.66 | **0.95 \***<br>**0.98 \***<br>**0.96 \***<br>**0.96 \*** | **1\*** |
| Out-Degree | mixed | Grooming | 0.89 | **sex(M)**<br>female age<br>male age<br>female rank<br>male rank | 0.87<br>**6.01**<br>1.54<br>4.29 | 1.07 | 0.54<br>**0.98\***<br>0.69<br>0.94 | **1\*** |
| In-Strength | mixed | Grooming | 0.67 | **sex(M)**<br>female age<br>male age<br>female rank<br>male rank | 1.06<br>7.63<br>0.83<br>9.7 | 0.52 | 0.19<br>0.8<br>0.19<br>0.89 | 0.84 |
| Out-Strength | mixed | Grooming | 0.73 | **sex(M)**<br>female age<br>male age<br>female rank<br>male rank | 1.81<br>1.04<br>1.2<br>2.81 | 1.72 | 0.6<br>0.44<br>0.54<br>0.77 | **1\*** |
| Local Transitivity | mixed | Grooming | 0.31 | **sex(M)**<br>female age<br>male age<br>female rank<br>male rank | 4.54<br>12.23<br>3.7<br>0.03 | 0.18 | **0.98\***<br>**1\***<br>**0.95\***<br>0.11 | **1\*** |
| Betweenness | mixed | Grooming | 0.56 | **sex(M)**<br>female age<br>male age<br>female rank<br>male rank | 3.55<br>1.91<br>2.1<br>2.76 | 1.48 | 0.68<br>0.5<br>0.55<br>0.63 | **1\*** |
| Eigenvector Centrality | mixed | Grooming | 0.84 | **sex**<br>female age<br>male age<br>female rank<br>male rank | 1.82<br>4.16<br>2.65<br>0.47 | 1.36 | 0.77<br>**0.95\***<br>0.89<br>0.42 | **1\*** |

**Table S4.** GAMM models for all integration measures in <u>same-sex grooming</u> networks. Significant effects in bold with *. DE = total model deviance explained. Significance of all smooth terms (age & rank) evaluated with observed F statistics compared to F statistics drawn from randomized networks.

| Response | Network sex | Behavior | DE | Predictors | F<sub>obs</sub> of smooths | % F<sub>obs</sub> > F<sub>ran</sub> |
| --- | --- | --- | --- | --- | --- | --- |
| In-Degree | same | Grooming | 0.09 | female age<br>female rank | 0.76<br>0.65 | 0.45<br>0.39 |
| In-Degree | same | Grooming | 0.38 | **male age**<br>male rank | 5<br>1.19 | **0.99\***<br>0.69 |
| Out-Degree | same | Grooming | 0.09 | female age<br>female rank | 0.76<br>0.65 | 0.24<br>0.2 |
| Out-Degree | same | Grooming | 0.54 | male age<br>male rank | 0.74<br>0.54 | 0.57<br>0.5 |
| In-Strength | same | Grooming | 0.5 | female age<br>female rank | 5.92<br>9.15 | 0.59<br>0.8 |
| In-Strength | same | Grooming | 0.64 | male age<br>male rank | 4.24<br>2.34 | 0.85<br>0.87 |
| Out-Strength | same | Grooming | 0.5 | female age<br>**female rank** | 5.92<br>9.15 | 0.93<br>**1\*** |
| Out-Strength | same | Grooming | 0.5 | male age<br>male rank | 1.55<br>2.14 | 0.51<br>0.9 |
| Local Transitivity | same | Grooming | 0.02 | female age<br>female rank | 1.14<br>1.46 | 0.61<br>0.67 |
| Local Transitivity | same | Grooming | 0.03 | male age<br>male rank | 0.34<br>0.6 | 0.39<br>0.53 |
| Betweenness | same | Grooming | 0.52 | female age<br>female rank | 1.9<br>1.84 | 0.27<br>0.23 |
| Betweenness | same | Grooming | 0.39 | male age<br>male rank | 3.03<br>0.54 | 0.43<br>0.16 |
| Eigenvector Centrality | same | Grooming | 0.42 | female age<br>female rank | 2.1<br>1.18 | 0.79<br>0.58 |
| Eigenvector Centrality | same | Grooming | 0.66 | **male age**<br>male rank | 6.07<br>0.08 | **0.98\***<br>0.16 |

**Table S5.** GAMM models for all SNA measures in <u>mixed-sex proximity</u> networks. Significant effects in bold with*. DE = total model deviance explained. Significance of the categorical variable sex evaluated with linear ß estimates, and all smooth terms (age & rank) evaluated with observed F statistics, each compared to ßs and F statistics drawn from randomized networks.

| Response | Network sex | Behavior | DE | Predictors | F<sub>obs</sub> of smooths | $\beta_{\text{obs}}$ of sex(M) | % F<sub>obs</sub> > F<sub>ran</sub> | % $\beta_{\text{obs}}$ > $\beta_{\text{ran}}$ |
| --- | --- | --- | --- | --- | --- | --- | --- | --- |
| Strength | mixed | Prox | 0.53 | **sex(M)** |  | 0.44 |  | **1\*** |
|  |  |  |  | female age | 3.49 |  | 0.94 |  |
|  |  |  |  | male age | 0.6 |  | 0.43 |  |
|  |  |  |  | female rank | 1.81 |  | 0.83 |  |
|  |  |  |  | male rank | 2.85 |  | 0.91 |  |
| Local Transitivity | mixed | Prox | 0.02 | **sex(M)** |  | 0 |  | 0.35 |
|  |  |  |  | female age | 0.2 |  | 0.25 |  |
|  |  |  |  | male age | 0.16 |  | 0.24 |  |
|  |  |  |  | female rank | 0.96 |  | 0.72 |  |
|  |  |  |  | male rank | 0.23 |  | 0.33 |  |
| Betweenness | mixed | Prox | 0.37 | **sex(M)** |  | -0.98 |  | 0.05 |
|  |  |  |  | female age | 4.61 |  | 0.78 |  |
|  |  |  |  | male age | 0.2 |  | 0.14 |  |
|  |  |  |  | female rank | 3.52 |  | 0.71 |  |
|  |  |  |  | male rank | 0.19 |  | 0.14 |  |
| Eigenvector Centrality | mixed | Prox | 0.71 | **sex(M)** |  | 0.5 |  | **1\*** |
|  |  |  |  | female age | 2.91 |  | 0.91 |  |
|  |  |  |  | male age | 2.35 |  | 0.87 |  |
|  |  |  |  | female rank | 1.26 |  | 0.68 |  |
|  |  |  |  | **male rank** | 21.25 |  | **1\*** |  |

**Table S6.**
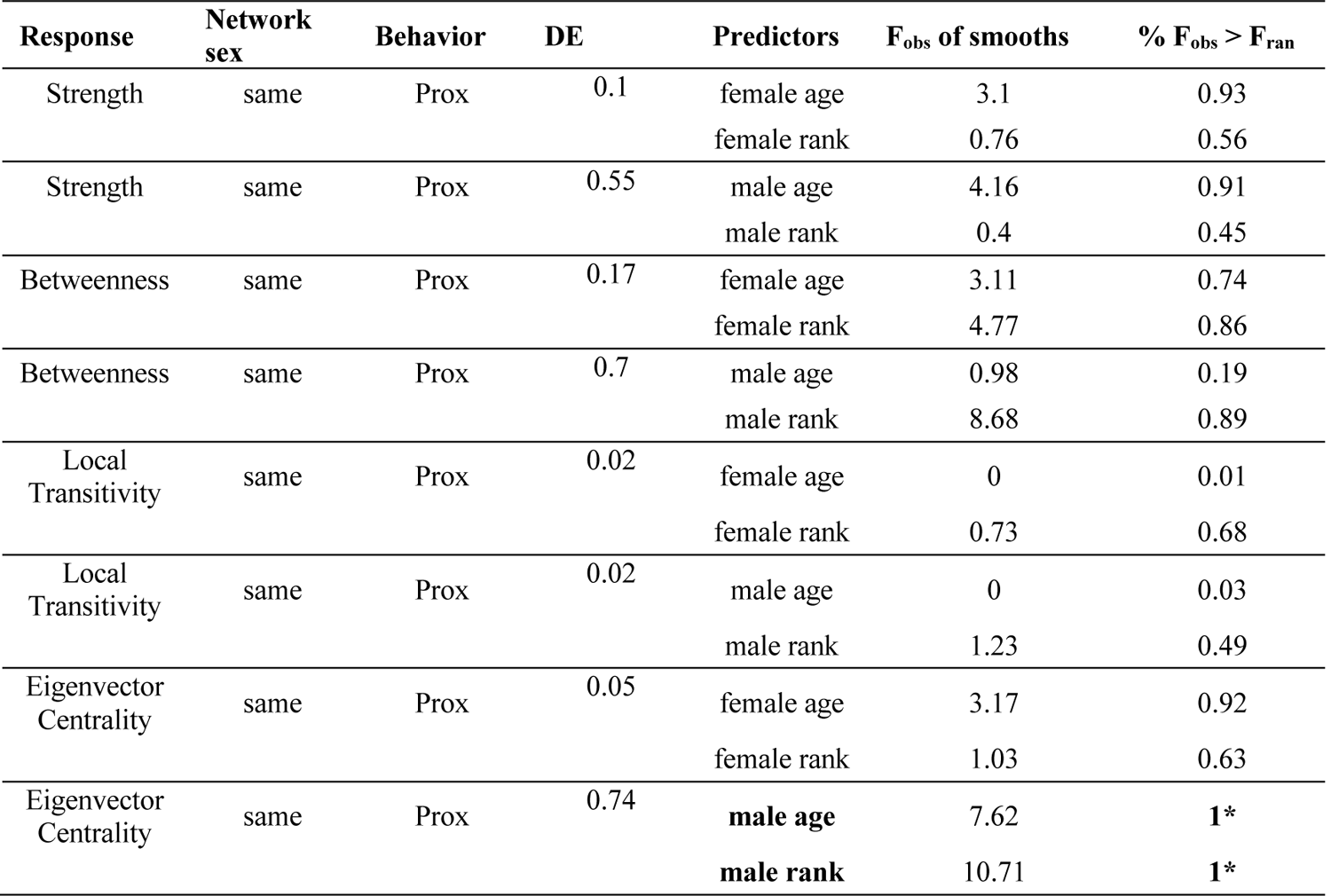
GAMM models for all integration measures in <u>same-sex proximity</u> networks. Significant effects in bold with *. DE = total model deviance explained. Significance of all smooth terms (age & rank) evaluated with observed F statistics compared to F statistics drawn from randomized networks.

**Table S7.**
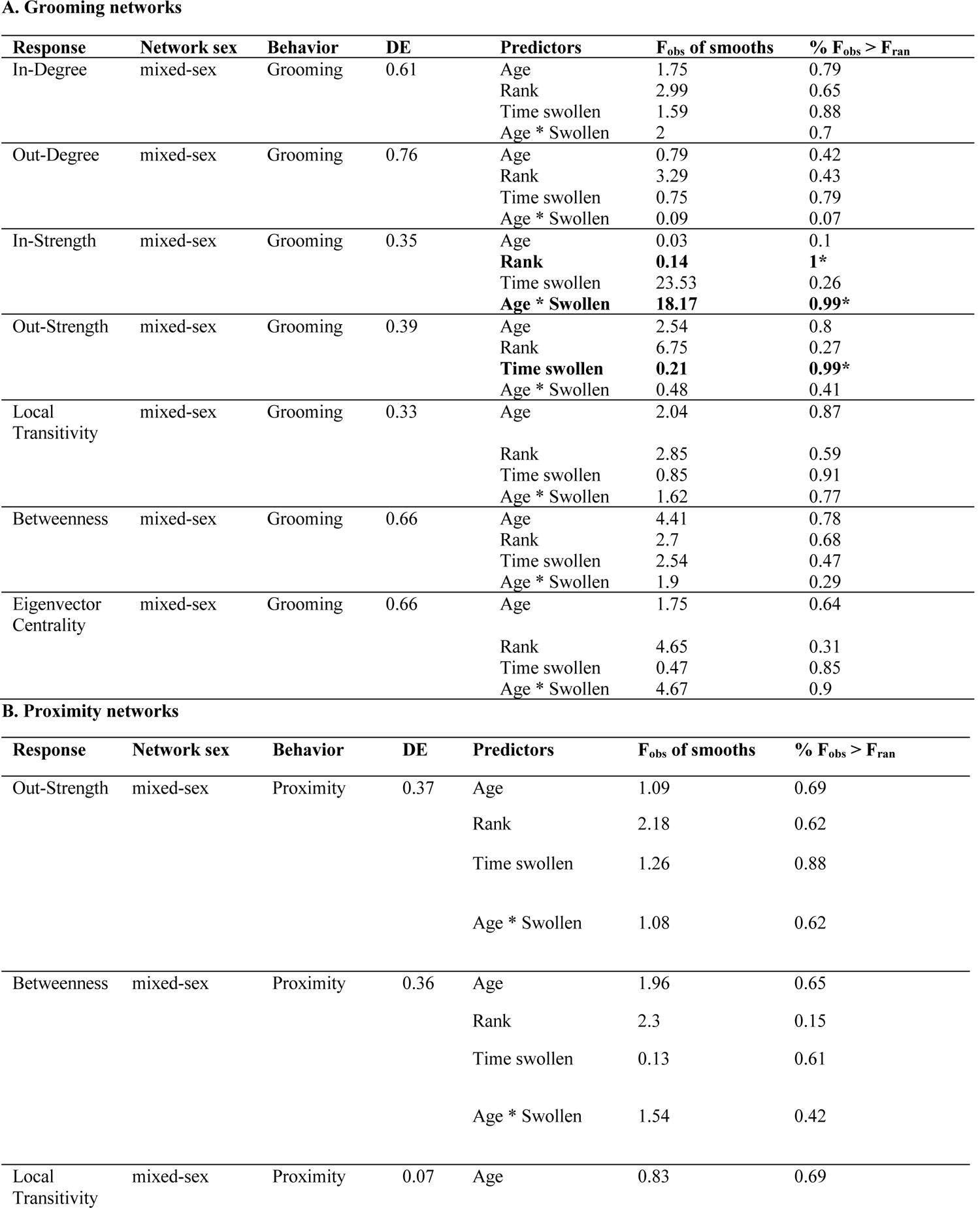
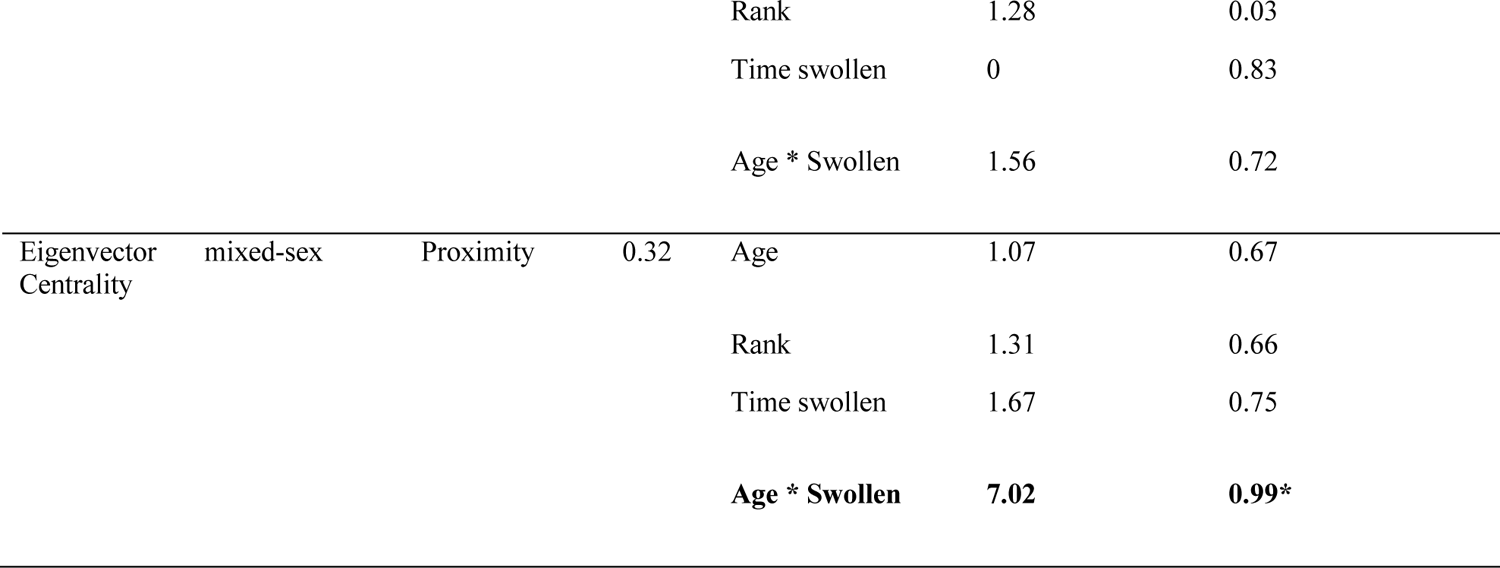
**Age effects independent of rank and time sexually swollen on female social integration in mixed-sex networks.** Significant effects in bold with*. DE = total model deviance explained. Significance of all smooth terms (age, rank, time swollen, and their interaction) evaluated with observed F statistics compared to F statistics drawn from randomized networks.

**Table S8.**
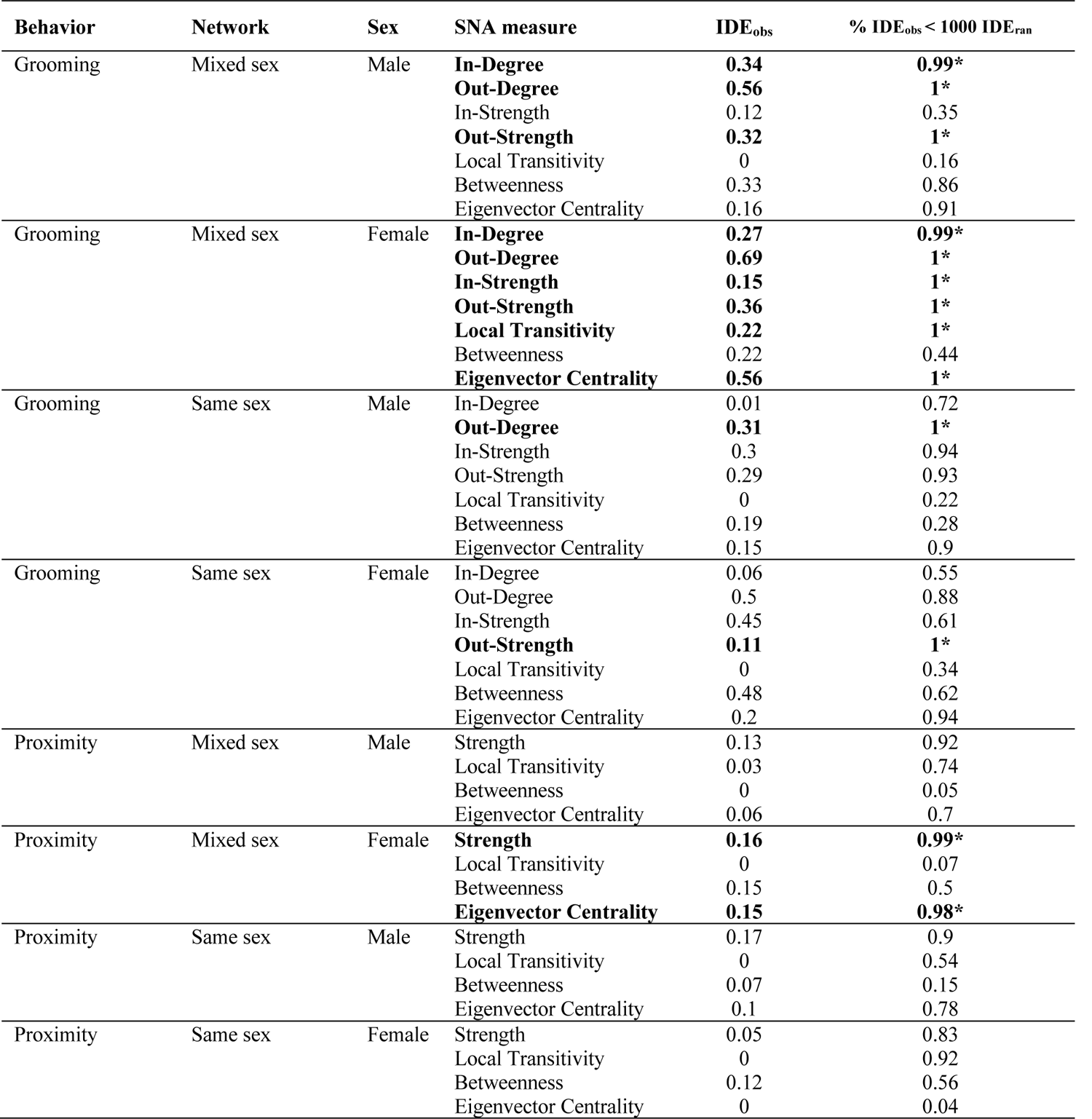
**Repeatability of integration measures by behavior, network type, and sex.** Repeatability statistic calculated by the observed deviance explained by individual ID alone (IDE_obs_) in Generalized Additive Mixed Models (GAMMs). Significance of IDE_obs_ evaluated by the proportion of 1000 deviances explained by ID in GAMMs on node-randomized data (IDE_ran_) that IDE_obs_ is less than.

**Table S9.**
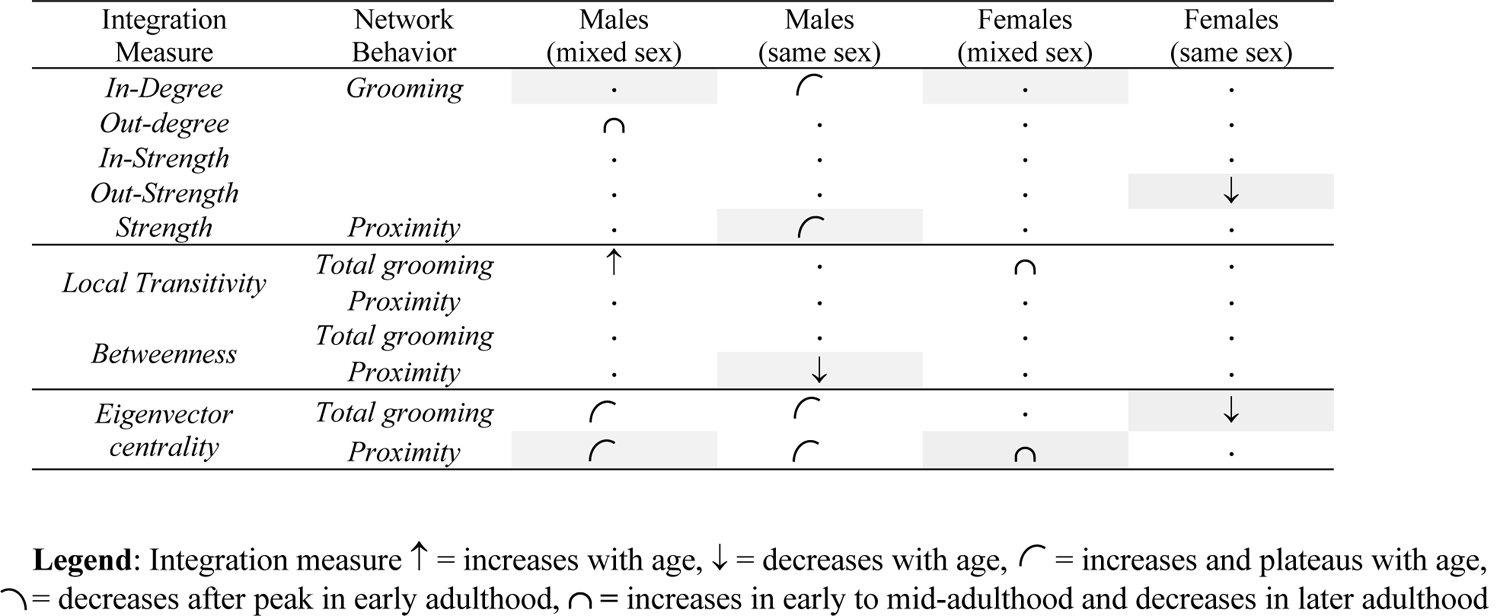
**Summary of age-alone model results:** Age-related changes in social network integration with shape or arrow describing any significant relationship between age and the given network measure. Effects are not controlling for dominance rank or time swollen. Shape and arrows describe significant relationships between age and a given network measure (see Legend; full model results in Tables S9-13). Dots indicate a non-significant pattern. Shading indicates a difference in significant patterns from rank-independent age models.

**Table S10.**
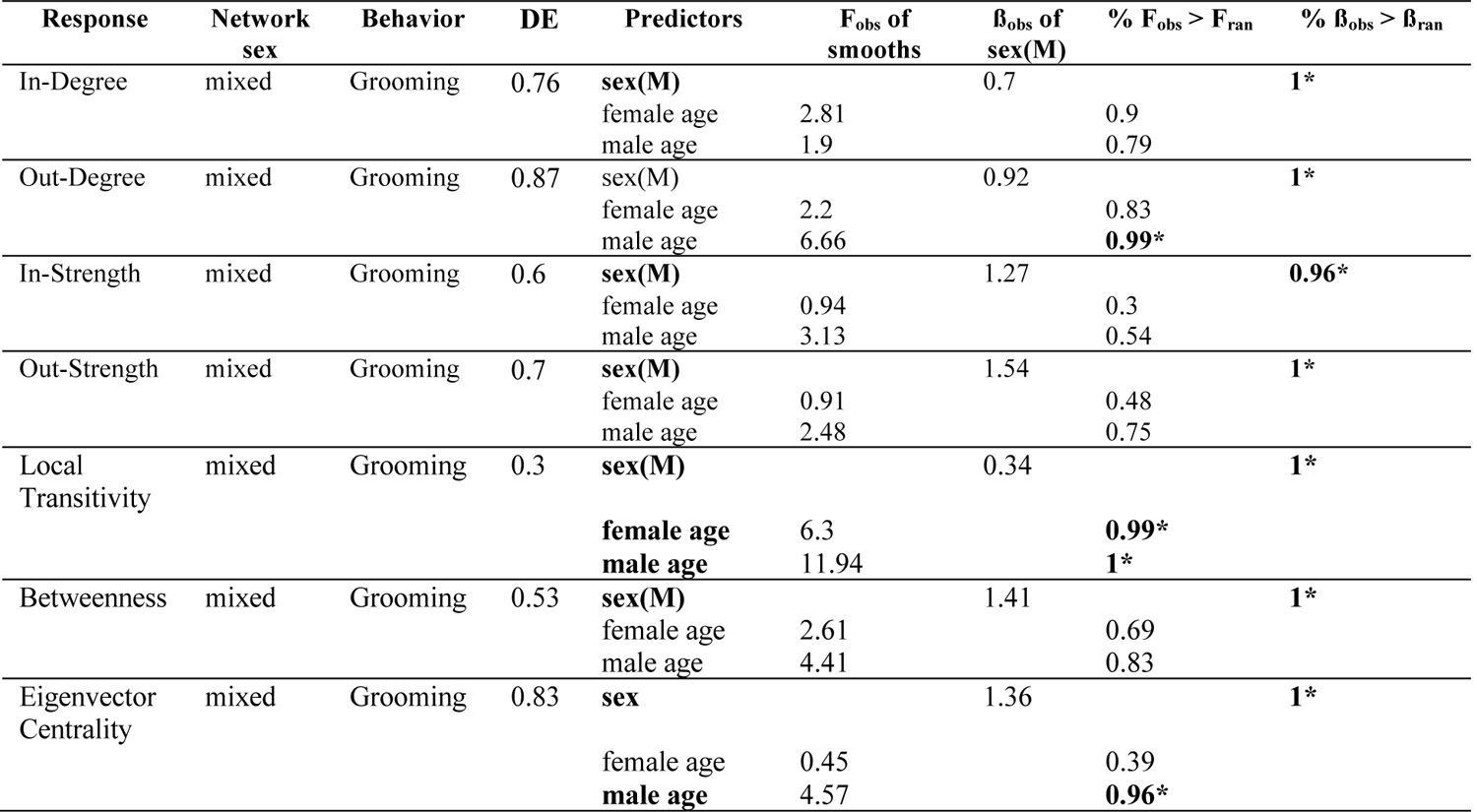
GAMM models with age alone as a predictor of integration measures in <u>mixed-sex grooming</u> networks. Significant effects in bold with *. DE = total model deviance explained. Significance of the categorical variable sex evaluated with linear ß estimates, and smooth term age evaluated with observed F statistics, each compared to ßs and F statistics drawn from randomized networks.

**Table S11.** GAMM models with age alone as a predictor of integration measures in <u>same-sex grooming</u> networks. Significant effects in bold with *. DE = total model deviance explained. Significance of smooth term age evaluated with observed F statistics compared to F statistics drawn from randomized networks.

| Response | Network sex | Behavior | DE | R | Predictors | F<sub>obs</sub> of smooths | % F<sub>obs</sub> > F<sub>ran</sub> |
| --- | --- | --- | --- | --- | --- | --- | --- |
| In-Degree | same | Grooming | 0.09 | 0.05 | female age | 0.46 | 0.38 |
| In-Degree | same | Grooming | 0.36 | 0.32 | **male age** | 7.1 | **1\*** |
| Out-Degree | same | Grooming | 0.66 | 0.6 | female age | 6.92 | 0.89 |
| Out-Degree | same | Grooming | 0.55 | 0.46 | male age | 0.96 | 0.66 |
| In-Strength | same | Grooming | 0.19 | 0.14 | female age | 0.85 | 0.35 |
| In-Strength | same | Grooming | 0.62 | 0.55 | male age | 3.51 | 0.81 |
| Out-Strength | same | Grooming | 0.35 | 0.25 | **female age** | 61.01 | **1\*** |
| Out-Strength | same | Grooming | 0.35 | 0.28 | male age | 2.24 | 0.68 |
| Local Transitivity | same | Grooming | 0.03 | 0.01 | female age | 0.46 | 0.42 |
| Local Transitivity | same | Grooming | 0.01 | 0 | male age | 0.77 | 0.57 |
| Betweenness | same | Grooming | 0.49 | 0.43 | female age | 4.36 | 0.63 |
| Betweenness | same | Grooming | 0.55 | 0.46 | male age | 0.96 | 0.3 |
| Eigenvector Centrality | same | Grooming | 0.42 | 0.36 | **female age** | 6.43 | **0.98\*** |
| Eigenvector Centrality | same | Grooming | 0.65 | 0.6 | **male age** | 7.04 | **0.99\*** |

**Table S12.**
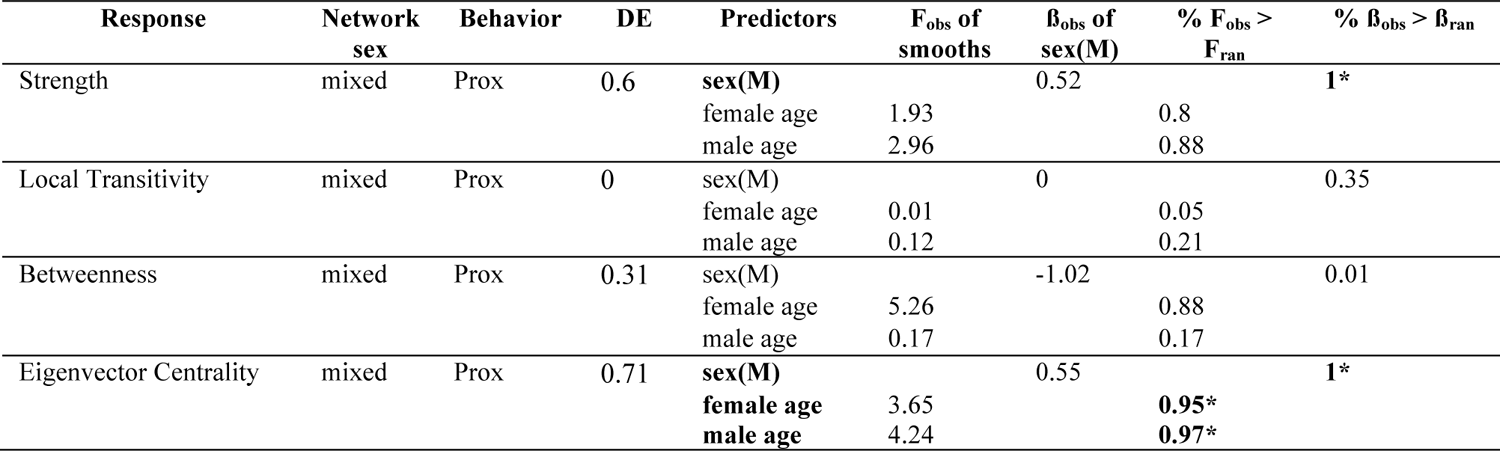
GAMM models with age alone as a predictor of integration measures in <u>mixed-sex proximity</u> networks. Significant effects in bold with*. DE = total model deviance explained. Significance of the categorical variable sex evaluated with linear ß estimates, and smooth term age evaluated with observed F statistics, each compared to ßs and F statistics drawn from randomized networks.

**Table S13.**
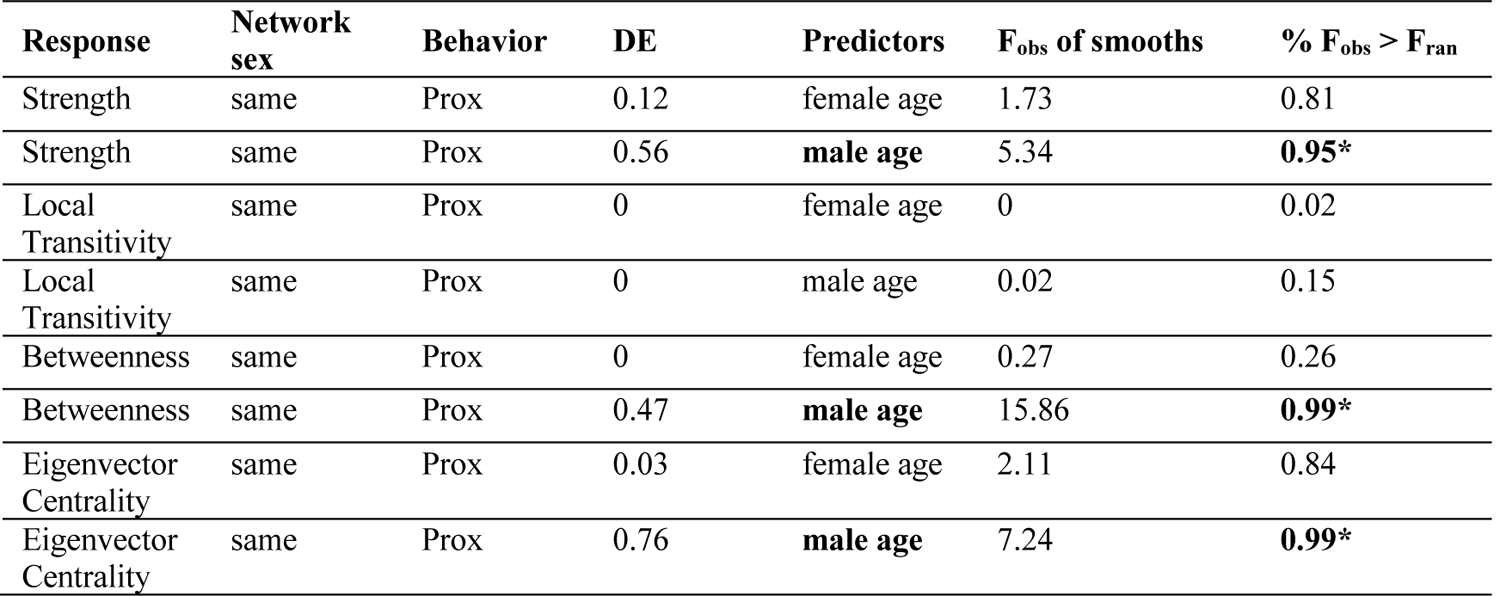
GAMM models with age alone as a predictor of integration measures in <u>same-sex proximity</u> networks. Significant effects in bold with *. DE = total model deviance explained. Significance of smooth term age evaluated with observed F statistics compared to F statistics drawn from randomized networks.

| Response | Network sex | Behavior | DE | Predictors | F<sub>obs</sub> of smooths | % F<sub>obs</sub> > F<sub>ran</sub> |
| --- | --- | --- | --- | --- | --- | --- |
| Strength | same | Prox | 0.12 | female age | 1.73 | 0.81 |
| Strength | same | Prox | 0.56 | **male age** | 5.34 | **0.95\*** |
| Local Transitivity | same | Prox | 0 | female age | 0 | 0.02 |
| Local Transitivity | same | Prox | 0 | male age | 0.02 | 0.15 |
| Betweenness | same | Prox | 0 | female age | 0.27 | 0.26 |
| Betweenness | same | Prox | 0.47 | **male age** | 15.86 | **0.99\*** |
| Eigenvector Centrality | same | Prox | 0.03 | female age | 2.11 | 0.84 |
| Eigenvector Centrality | same | Prox | 0.76 | **male age** | 7.24 | **0.99\*** |

## Figure captions

**Figure 1.** Age ranges of observation for each study subject (22 F & 16 M; 122 female-years, 78 male-years). Focal observations were continuous over each age window.

**Figure 2.** Social integration measures by age in mixed and same-sex grooming networks. Male data represented by blue triangles and blue dashed GAM smooth, female data represented by red circles and red solid GAM smooth. Smooths are conditional effects of age on social integration, controlling for rank, created using the R functions visreg and mgcv::gam within ggplot2.

**Figure S1.**
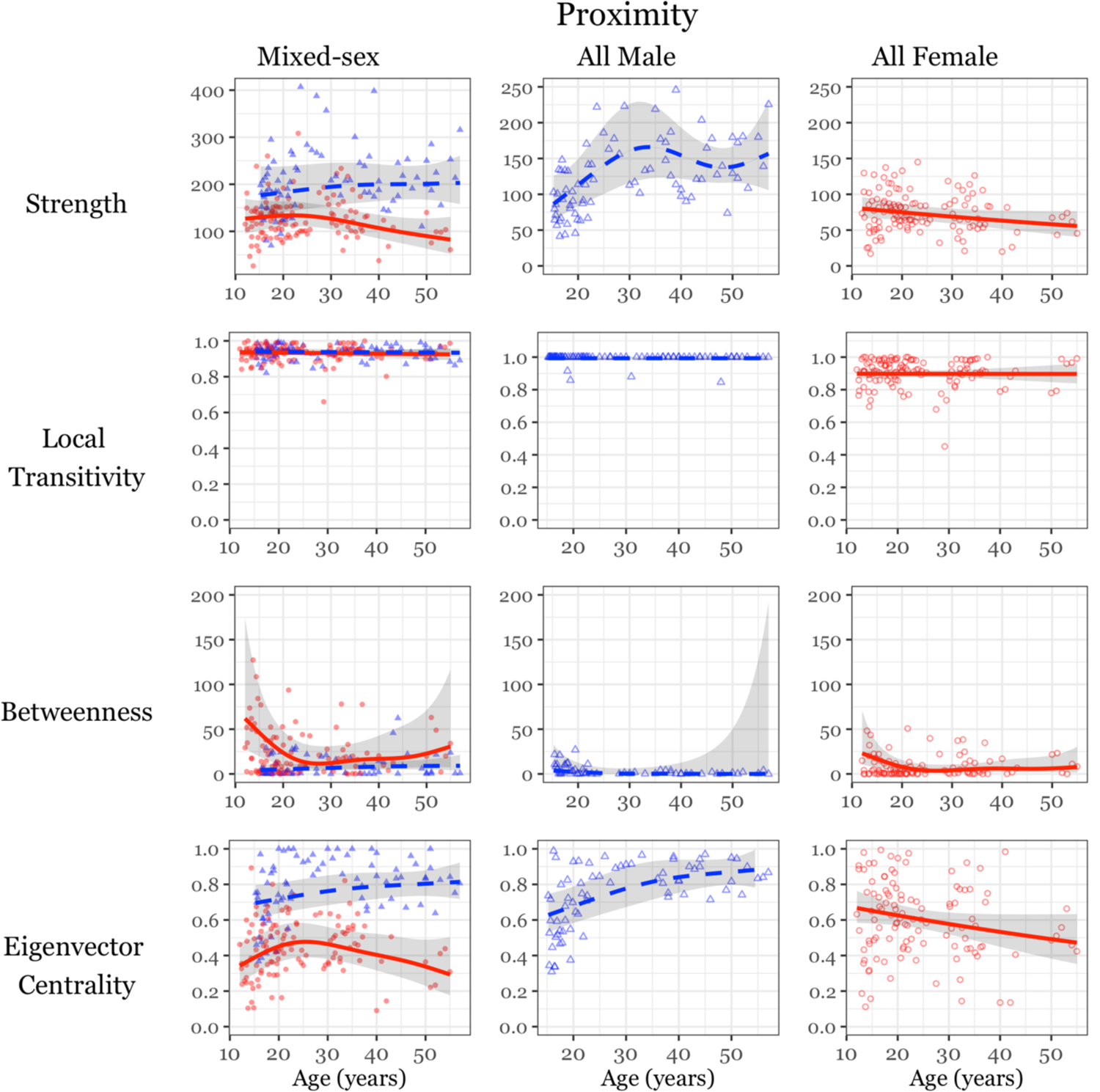
All social integration measures by age in mixed and same-sex **<u>proximity</u>** networks. Male data represented by blue triangles and blue dashed GAM smooth, female data represented by red circles and red solid GAM smooth. Smooths are conditional effects of age on social integration, controlling for rank, created using the R functions visreg and mgcv::gam within ggplot2.

**Figure S2.**
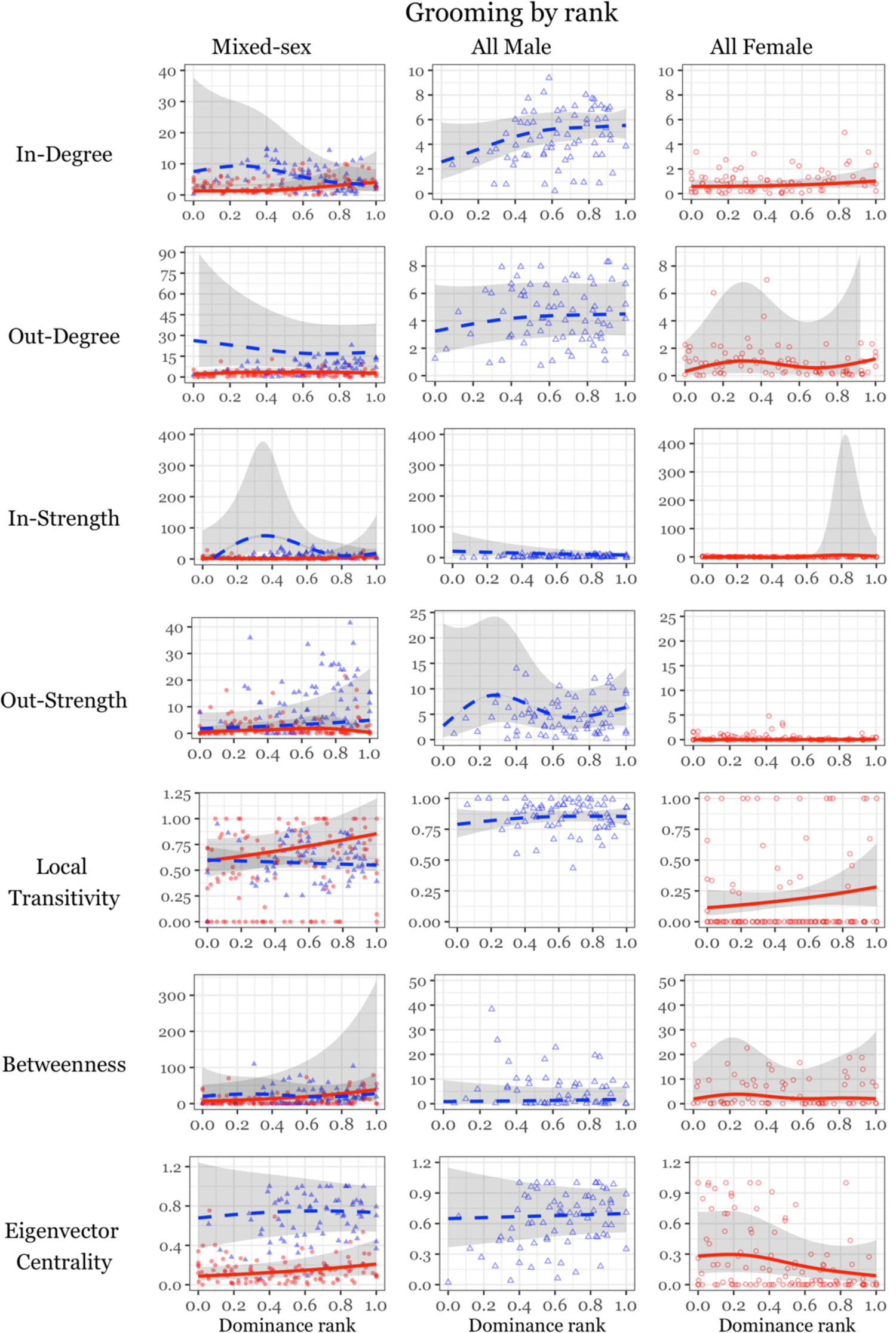
Social integration in mixed and same-sex grooming networks by **<u>dominance rank</u>**. Male data represented by blue triangles and blue dashed GAM smooth, female data represented by red circles and red solid GAM smooth. Smooths are conditional effects of age on social integration, controlling for rank, created using the R functions visreg and mgcv::gam within ggplot2.

**Figure S3.**
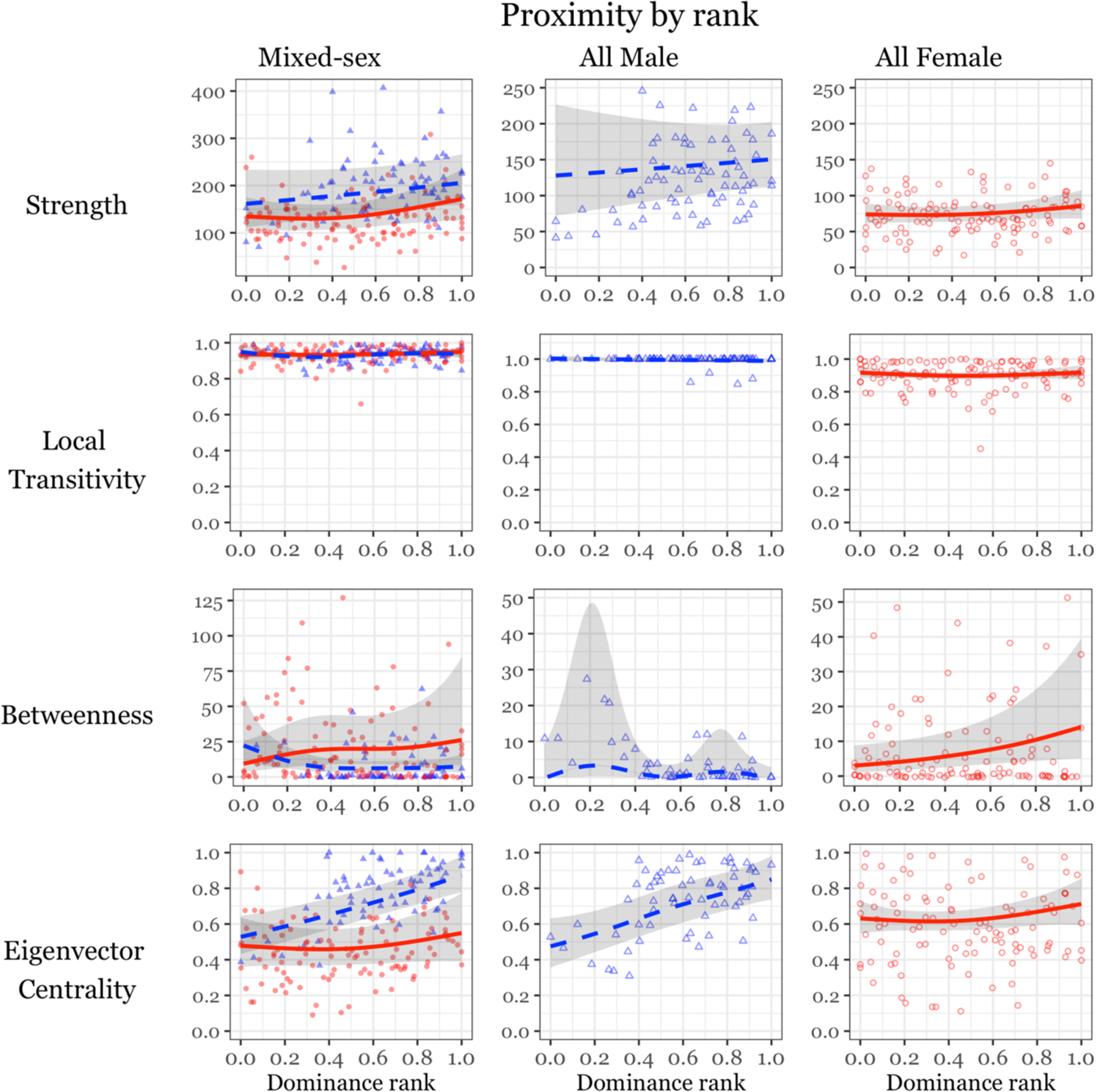
Social integration in proximity networks by **<u>dominance rank</u>**. Male data represented by blue triangles and blue dashed GAM smooth, female data represented by red circles and red solid GAM smooth. Smooths are conditional effects of age on social integration, controlling for rank, created using the R functions visreg and mgcv::gam within ggplot2.

**Figure S4.**
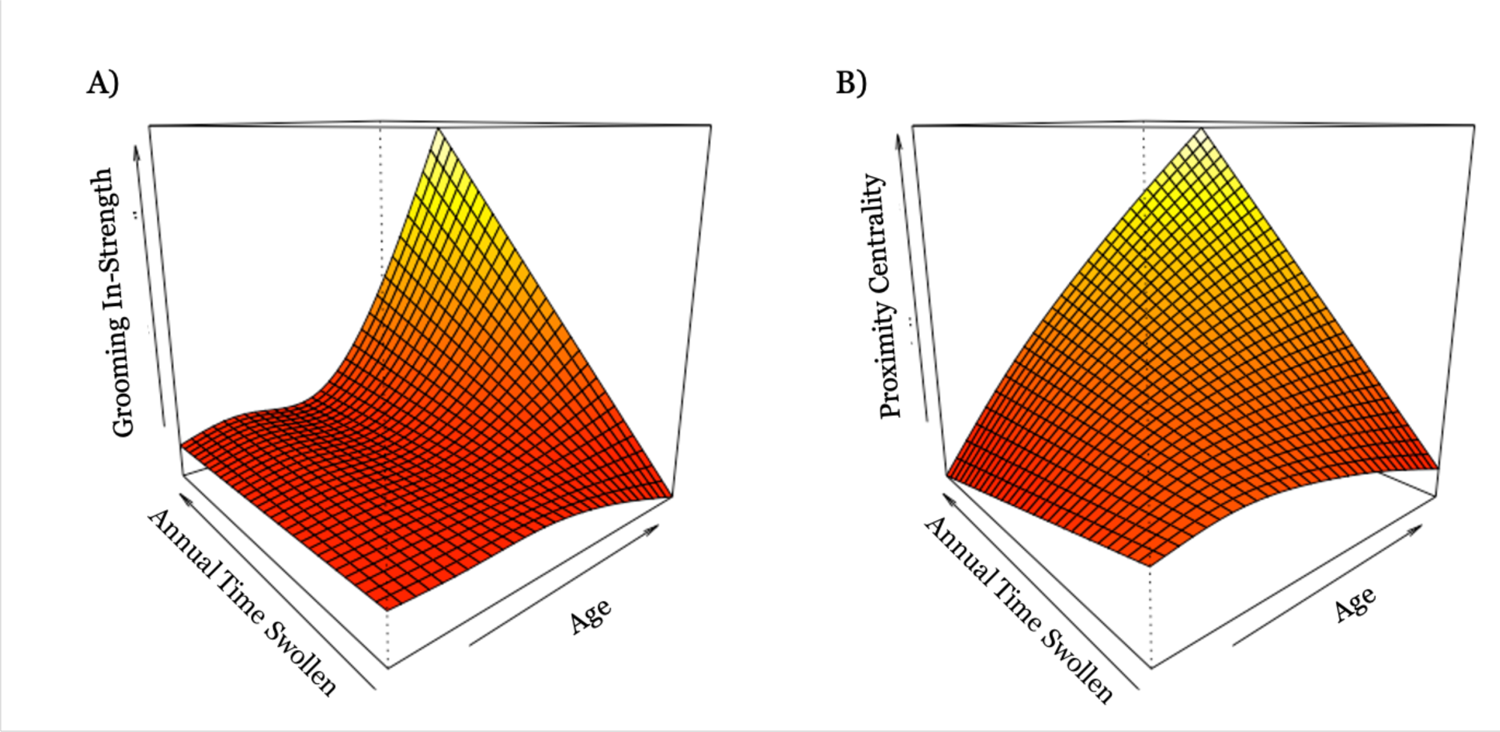
Changes in female A) grooming in-strength and B) proximity centrality in mixed sex networks as a product of age and annual time fully swollen. Plots created using the vis.gam function in R’s mgcv package.

